# Ion channel and receptor-mediated regulation of axonal conduction reliability in sympathetic preganglionic neurons

**DOI:** 10.64898/2026.06.03.729634

**Authors:** Mallika Halder, Shawn Hochman

**Author notes:** Competing Interests: The authors declare that they have no competing interests.

## Abstract

Sympathetic preganglionic neurons (SPNs) provide the sole spinal output to the peripheral sympathetic nervous system. Although sympathetic control is traditionally attributed to synaptic integration within the spinal cord and ganglia, the reliability of spike propagation along SPN axons themselves has received little attention. Here, and in companion papers, we show that axonal conduction in adult mouse thoracic SPNs is highly modifiable and constitutes a critical site of sympathetic gain control. Using an *ex vivo* preparation preserving intact paravertebral and splanchnic pathways while blocking synaptic transmission, we recorded compound action potentials evoked across multiple ganglia. Slower-conducting, unmyelinated SPN axons, particularly those with branching axons traversing the interganglionic nerve (IGN), exhibited pronounced, temperature-dependent conduction failures. Elevation of temperature produced membrane hyperpolarization and loss of conduction, consistent with activation of temperature-sensitive K_2_P leak channels, as supported by pharmacological evidence. Pharmacological activation of TREK-family channels with riluzole or arachidonic acid preferentially suppressed conduction in these axons. In contrast, blockade of voltage-gated K^+^ channels with 4-aminopyridine (4-AP) robustly facilitated conduction, recruited previously silent axons, and restored propagation under conditions of temperature-induced failure. Surprisingly, tetraethylammonium (TEA) block of K^+^ channels were without effect or depressant. Transmitter systems further shaped axonal reliability: agonists and antagonists of GABA_A_ receptors, as well as cholinergic manipulations, selectively depressed conduction in slow, branching axons. Together, these findings establish SPN axons, particularly slow-conducting branching fibers, as an active and dynamically regulated substrate for sympathetic output control, revealing a presynaptic mechanism with implications for autonomic physiology and disease.

**SIGNIFICANCE:** Sympathetic output is commonly viewed as being regulated primarily through synaptic integration within spinal and autonomic circuits, while axons are often treated as passive transmission elements. Emerging evidence suggests this assumption is incomplete, particularly in slowly conducting and highly branched sympathetic preganglionic neuron (SPN) axons that may operate near the limits of conduction reliability. This study identifies branch point conduction as a dynamic and pharmacologically modifiable control mechanism governing sympathetic signal transmission. By demonstrating selective vulnerability of distinct SPN populations and revealing strong modulation by potassium channel mechanisms, these findings establish axonal conduction security as an underappreciated site of autonomic gain control. These mechanisms may represent novel therapeutic targets for restoring autonomic function after spinal cord injury and related disorders.

## INTRODUCTION

Sympathetic preganglionic neurons (SPNs) form the final common output of the central nervous system to the peripheral sympathetic network. These neurons integrate descending and local spinal signals and transmit commands to postganglionic neurons innervating cardiovascular, thermoregulatory, metabolic, and visceral effector organs. Thoracic SPNs exhibit a distinctive axonal organization dominated by unmyelinated axons(Halder and Hochman, 2026; Halder et al., 2026). Many SPNs project to prevertebral ganglia via the unbranching splanchnic nerve; others project through branching pathways within the paravertebral sympathetic chain to act on postganglionic neurons. These unmyelinated axons branch extensively to generate anatomical divergence to enable individual SPNs to influence large populations of postganglionic neurons. It also introduces potential points of vulnerability for spike propagation.

Classically, regulation of sympathetic output has been viewed primarily through the lens of synaptic integration and neurotransmitter release within sympathetic ganglia. In contrast, the axon itself has often been treated as a passive conduit for reliable spike transmission (McLachlan, 2003). However, studies in both central and peripheral nervous systems increasingly demonstrate that axonal conduction, particularly in small-diameter unmyelinated fibers, is neither invariant nor guaranteed. Action potential propagation can fail at branch points, during high-frequency activity, or under altered metabolic and thermal conditions (Bostock et al., 1981; Wall, 1995; Pekala et al., 2016). Such failures can profoundly shape population output without altering presynaptic firing rates.

Our recent work in mice has established that SPN axons encompass a broad spectrum of conduction velocities and that the slowest-conducting unmyelinated axons show the greatest variability in recruitment across repeated stimuli, consistent with intermittent conduction failure (Halder et al., 2026). These failures are particularly prominent in axons presumed to branch within the sympathetic chain, suggesting that axonal geometry is a critical determinant of reliability. Moreover, increasing temperature within the physiological range strongly depresses conduction in these axons, indicating that sympathetic output may be gated by biophysical mechanisms intrinsic to the axon (Halder et al., 2026).

Ion channels expressed along axons are well positioned to regulate spike propagation. Voltage-gated potassium (K_V_) channels shape action potential repolarization and can limit propagation in axons (Debanne et al., 1997; Debanne et al., 2011), while two-pore domain potassium (K_2_P) leak channels could also regulate propagation by regulating resting membrane potential and membrane resistance (Enyedi and Czirják, 2010). In addition, axonal receptors traditionally associated with synaptic transmission, including GABA_A_, serotonergic, and cholinergic receptors, can modulate excitability through tonic or constitutive activity (Trigo et al., 2008; Hari et al., 2022). Transcriptomic analyses reveal that SPNs express a diverse complement of these channels and ionotropic receptors (Alkaslasi et al., 2021; Blum et al., 2021), raising the possibility that sympathetic output is regulated at the level of axonal conduction itself.

Here we test the hypothesis that conduction along thoracic SPN axons is dynamically modulated by ion channels. Using an ex vivo adult mouse preparation that preserves intact sympathetic pathways while eliminating synaptic transmission, we examine population spike propagation in branching axons of the interganglionic nerve (IGN) and unbranching axons of the splanchnic nerve. By combining electrical and optogenetic recruitment with targeted pharmacology, we identify mechanisms that selectively gate conduction in vulnerable axonal populations and define a previously underappreciated mode of sympathetic gain control.

## METHODS

### Multisegmental ex vivo paravertebral preparation

An overview of the anatomical organization is provided in **Figure 1A**. Experiments were undertaken in adult choline acetyltransferase (ChAT)-channelrhodopsin (ChR2) mice: driver line is ChAT-IRES-cre (JAX: 006410); reporter line is R26-ChR2-eYFP (JAX: 012569). Details of the dissection have been provided (Halder et al., 2021). Briefly, adult (8 weeks+) mice of both sexes were used. No sex specific differences were observed. Mice were euthanized with intraperitoneal injection of urethane (2g/kg). The thoracic vertebral column and adjacent ribs weree excised and transferred to a dish containing ice cold, oxygenated (95% O_2_ / 5% CO_2_) high-Mg^2+^/low-Ca^2+^ solution containing (in mM), [NaCl 128, KCl 1.9, MgSO_4_ 13.3, CaCl_2_ 1.1, KH_2_PO_4_ 1.2, glucose 10, NaHCO_3_ 26]. Following a complete laminectomy and vertebrectomy, the spinal cord (SC) and dorsal roots (DR) were exposed and removed. The remaining thoracic chain ganglia, in continuity with communicating rami, spinal nerves and ventral roots (**Fig. 1A_2_**) weree cleaned of excess fat and muscle. The tissue was transferred to a Sylgard 170 (Dow Inc.) silicone-bottomed recording chamber and superfused at ^~^40ml/minute with oxygenated extracellular solution maintained at pH of 7.4 and containing (in mM), [NaCl 128, KCl 1.9, MgSO_4_ 1.3, CaCl_2_ 2.4, KH_2_PO_4_ 1.2, glucose 10, NaHCO_3_ 26], and allowed to rest for 1 hour.

**Figure 1.**
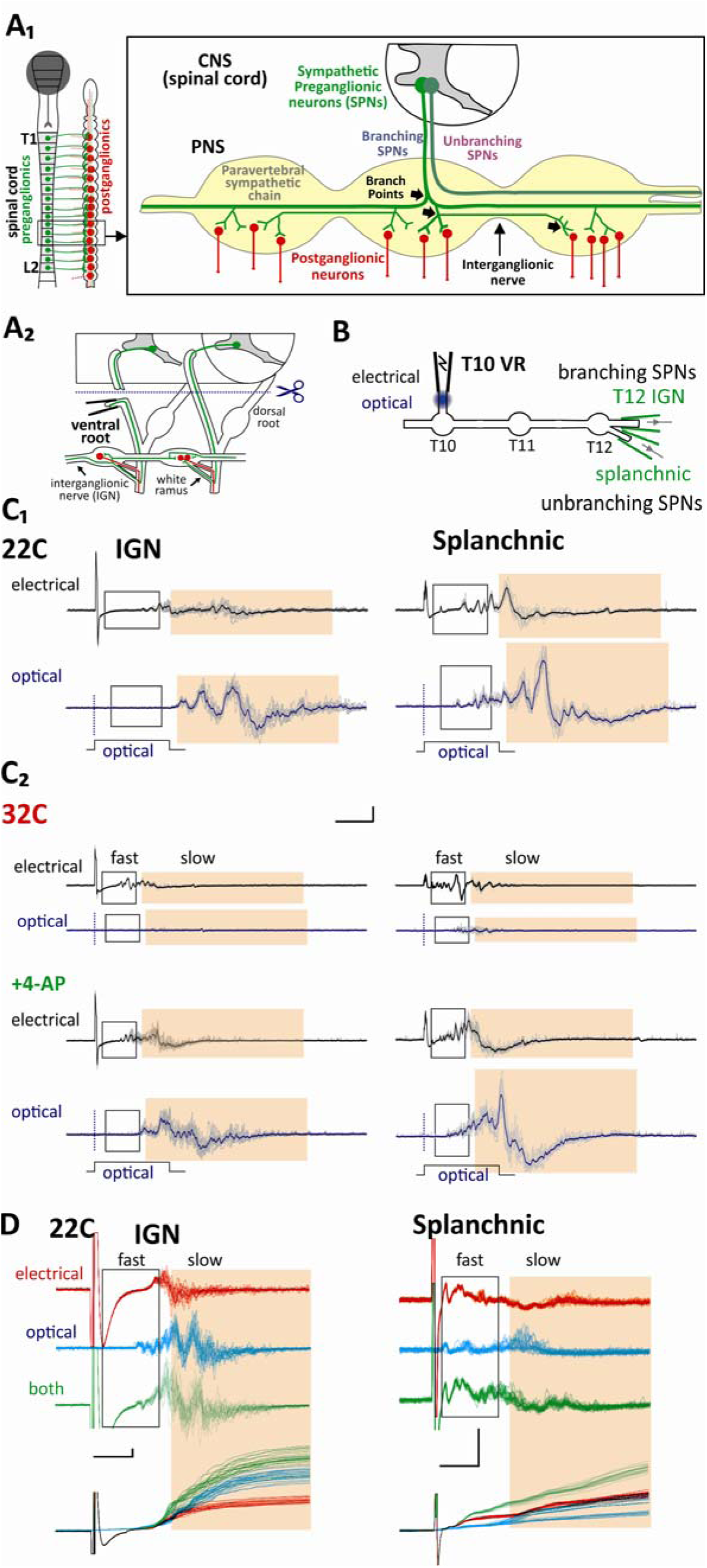
Anatomical organization and experimental approach for quantification of fast and slow conducing axonal population in branching (IGN) and unbranching (splanchnic) SPN axons. [A_1_]. All sympathetic preganglionic neuron (SPN) cell bodies are in the spinal cord. Axons of individual SPNs branch to innervate several postganglionic neurons in several paravertebral chain ganglia. Other SPNs do not innervate chain ganglia and travel unbranched along the interganglionic nerve before exiting to innervate prevertebral ganglia located nearer to target organs (not shown). **[A_2_]** Approach enabling selective activation of preganglionics via stimulation of the ventral root. **[B]** Experimental setup and the recording of CAPs from SPN axonal projections at T12 IGN and splanchnic nerve following electrical or optogenetic recruitment of axons from the T10 ventral root. **[C]** Conduction time ranges used to divide axonal populations into fast and slow conducting populations at 22C (C_1_) and 32C (C_2_). Overlaid raw records of 10 episodes recorded at 1/60 seconds illustrate temperature-, nerve- and stimulus method-dependent differences in axonal populations recorded (thick traces are overlaid averages). Boxed region identifies faster while orange shaded region identifies slower conducting axons for separate quantification in both IGN and splanchnic nerves. Note that at 32C, there is limited capacity for axonal conduction by optical stimuli, but this is unmasked by administration of 4-AP (100 μM). Optical stimulus onset time is shown by a vertical dotted line. **[D]** Electrical, optical, and simultaneous electrical and optical evoked responses support recruitment of at least partly distinct populations. Top. Raw records. Bottom. Rectified integrals of raw recordings overlaid to highlight differences in population responses. All records are high-pass filtered at 100 Hz. Scale bars for C and D are 200μV, 10ms and 20μV, 10ms, respectively.

### Electrophysiology

The ganglia of the paravertebral chain was made visible and glass suction electrodes are placed delicately to ensure stable recordings for extended periods. Stimulation of the sympathetic preganglionic neuron (SPN) axons was undertaken from T10 ventral roots (**Fig. 1B**). A glass suction electrode was positioned for stimulation (200-250 *µ*m tip diameter) while glass suction recordings electrodes placed on the caudal cut ends of T12 interganglionic nerve (IGN) and splanchnic nerve using fabricated ‘trumpet’ shaped electrodes to generate tight seals (minimal internal diameter 100-125 *µ*m) (Halder et al., 2021). All recorded data were digitized at 50 kHz (Digidata 1322A 16 Bit DAQ, Molecular Devices, U.S.A.) with pClamp acquisition software (v. 10.7 Molecular Devices). Recorded signals were amplified (5000x) and low pass filtered at 3 kHz using in-house amplifiers. As recordings of population axonal compound action potential (CAP) events could include synaptically-mediated postganglionic neuronal spiking responses, we blocked synaptic activity with the nicotinic ganglionic acetylcholine (ACh) receptor antagonist hexamethonium (100 μM) (Mason, 1962) and included pancuronium (20 µM) to prevent possible intercostal muscle activity (both from Sigma-Aldrich). Unless otherwise stated, experiments were undertaken at 32°C. This temperature was chosen instead of 36°C because in preliminary experiments we determined that 32°C increased duration of preparation viability to >6 hours, which was necessary for prolonged pharmacological studies.

### Electrical recruitment of cholinergic axons in ventral roots

Electrical stimulation threshold was found to be ∼50µA, 50µs in the T12 IGN and splanchnic nerve recordings in most preparations with maximal recruitment typically achieved at 200µA, 200µs stimulation (n=5). To ensure supramaximal recruitment of slow-conducting axons, electrical stimulation intensity for experiments was kept at 200µA, 500µs, as longer pulse durations ensure recruitment of unmyelinated axons (Thompson et al., 1990; Brocker and Grill, 2013).

### Optical recruitment of cholinergic axons in ventral roots

The use of ChAT-CHR2 mice enabled comparison of electrically evoked axonal responses in ventral roots to selective optogenetic recruitment of cholinergic axons. This is because thoracic ventral roots are also thought to contain ventral root afferents (Coggeshall et al., 1977). Optogenetic stimulation was performed using custom-built calibrated Clampex-controlled laser diode boxes. ThorLabs M61L01 fiber optic cables (105 µm, 0.22 NA) were placed to target the same ventral root used in all experiments. Recruitment profiles were first compared at different optical intensities and durations. Responses could not be evoked by optical pulses shorter than 2ms with the fastest conducting unit observed with a 5 ms pulse and peak responses seen with 10 or 20ms pulses. In 4/5 experiments, a duration of 20 ms optical stimulation was needed to maximally recruit SPN populations. Maximal recruitment was usually achieved at an optical intensity of ∼6.3mW/mm^2^. Consequently, 20 ms pulse durations and 6.6mW/mm^2^ were used to ensure supramaximal recruitment. To enable separate analysis of fast and slow conducting SPN axonal subpopulations, we identified these populations based on alignment of conduction times of characteristic electrically evoked responses. This generally aligned the onset of recruitment ∼5 ms into the optical pulse.

### Analysis

Electrically and optically evoked SPN compound action potentials (CAPs), were recorded following VR stimulation once every 60 seconds. Recorded CAPs included events that span a large range of conduction velocities which are also temperature-dependent **(Table 1**)(Halder and Hochman, 2026). Populations were divided into fast and slow CV-subpopulations depending on latency of arrival from time of stimulation (**Table 2**). For each CV population, 10 episodes were captured for each condition then rectified and integrated using Clampfit software to quantify responses as total area (mV.ms). Unless otherwise stated all analysis was undertaken following high pass filtering (100Hz single pole RC). At this point background noise was accounted for by subtracting an equal duration of baseline noise to provide a singular, comprehensive measure of CAP size. All 10 episodes were then averaged to ensure a more accurate representation of the CAP. Ideally, comparisons across preparations would involve recordings at the same site and use the same electrodes consistently. However, in most experimental series, variability in evoked responses across preparations was large, so observed changes in individual experiments are presented as normalized to their baseline magnitude responses.

**Table 1.** Conduction velocity range (in m/s) of electrically-evoked thoracic SPN axons at 22° and 32°C (n=7, except optical IGN [n=6]).

| Electrical | 22°C (m/s) | 32°C (m/s) |
| --- | --- | --- |
| Interganglionic Nerve | 0.10±0.01- 0.74±0.25 | 0.18±0.01 - 1.40±0.51 |
| Splanchnic Nerve | 0.11±0.01 - 1.76±0.48 | 0.18±0.02 - 2.86±0.52 |
| Optical |  |  |
| Interganglionic Nerve | 0.09±0.004 - 0.95±0.44 | 0.21±0.03 - 1.27±0.41 |
| Splanchnic Nerve | 0.10±0.01– 5.40±1.59 | 0.23±0.04 - 3.14±1.03 |

**Table 2.** Conduction time ranges from start of stimulus used to quantify magnitude changes in fast and slow conducting axons at 22° and 32°C.

| T° & CV | Electrical/Optical start time; epoch duration |
| --- | --- |
| 22°C fast | 2.5/4.5 ms from stim. onset; 14.5ms duration |
| 22°C slow | 20/22ms from stim. onset; 43.1ms duration |
| 32°C fast | 2/3 ms from stim. onset; 9ms duration |
| 32°C slow | 12.5/13.5ms from stim. onset; 43.1ms duration |

### Pharmacology

Drugs, which were stored in 0.1, 1, 10 or 10mM stock solutions at −20◦C, were added to the reservoir of a perfusion bath (typically 100mL) with drug equilibration achieved via rapid recirculation. Drugs were applied by bath superfusion for between 5 and 30min until a maximum response was observed. Chemicals were obtained from Sigma Aldrich. L655,708, picrotoxin, bicuculline, muscimol, acetylcholine, neostigmine arachidonic acid, riluzole, 4-AP, ω-conotoxin GVIA, and ω-agatoxin IVA were added in concentrations previously reported in vitro (Alkadhi, 2008; Currò, 2010; Ivanov and Zilberter, 2011; Shreckengost et al., 2020; Bryson et al., 2025).

### Statistics

When a single drug was added, paired t-tests were used to compare baseline to drug condition. When more than one drug was used before washout, repeated measures ANOVA was used to determine significant effects by comparing to baseline control values. For post-hoc detailed pairwise comparisons, a Bonferroni t-test was utilized. Values are represented as mean percent of baseline ± standard error (SEM).

## RESULTS

The range of conduction velocities recorded following supramaximal electrical and optical stimulation is shown in **Table 1**. Mean slowest and fastest conduction velocities were comparable though optical tended to recruit additional faster conducting units. The conduction time ranges used to divide axonal populations into fast and slow conducting populations is shown in **Table 2** at both room temperature (22°C) and 32°C as well as for characterization following electrical versus optogenetic stimulation. Example recordings depict evoked responses in their separation into fast and slow categories for both splanchnic and IGN nerve at both temperatures (**Fig. 1C**). Comparison between electrically and optically evoked responses identify differences in recruitment. First, a surprising observation was that conduction of optically-recruited axons was particularly sensitive to conduction failure at higher temperature (32°C; **Fig. 1C_2_**). As shown, conduction failures could be restored pharmacologically with 4-AP. The effects of 4-AP and other modulatory actions on conduction are explored in detail in the sections below. Second, the combined sum of simultaneously delivered electrical and optical stimuli is greater than each recruited individually, indicating partial recruitment of independent axonal populations (Table 3; **Fig. 1D**). These differences will be compared in detail in studies below where relevant.

**Table 3.** Comparing the amplitude of electrical, optical, versus co-stimulation evoked responses from the same ventral root. Values represent % amplitude of the rectified integral response normalized to mean values evoked by electrical stimulation in T12 IGN and splanchnic nerve recordings (n=3).

|  | electrical | optical | combined |
| --- | --- | --- | --- |
| IGN | 100±30 | 139±58 | 241±58 |
| splanchnic | 100±50 | 158±26 | 266±77 |

### SPN conduction block with broad-spectrum GABA_A_R antagonists

Postganglionic neurons do not express GABA_A_Rs (Furlan et al., 2016) so all actions must occur pre-synaptically on SPNs. To investigate actions of modulators of GABA_A_Rs, we sequentially applied GABA_A_R blockers at 32°C. Examples responses are shown in **Figure 2A**. For slower-conducting IGN axons there was a trend toward progressive reduction in response with each blocker applied. The α_5_-GABA_A_R GABA binding site antagonist (inverse agonist) L655,708 (20μM) did not affect SPN axons consistent with limited expression reported for α_5_(Wang et al., 2008; Alkaslasi et al., 2021). Subsequent applications of the open channel non-competitive antagonist picrotoxin (50μM) (Newland and Cull-Candy, 1992) and competitive GABA_A_R antagonist bicuculline ultimately depressed slower-conducting IGN responses to 42.4±8.0% of baseline values (p<0.01, n=6, **Fig. 2B**). Faster-conducting IGN axons were not depressed by these GABA_A_R antagonists (post hoc t-testing), but overall population differences emerged (p<0.01; RM ANOVA) due to significant reduction after drug washout (55.9±16.2% of baseline values; p<0.01). Splanchnic fast and slow axon populations were not affected by GABA_A_R blockers.

**Figure 2.**
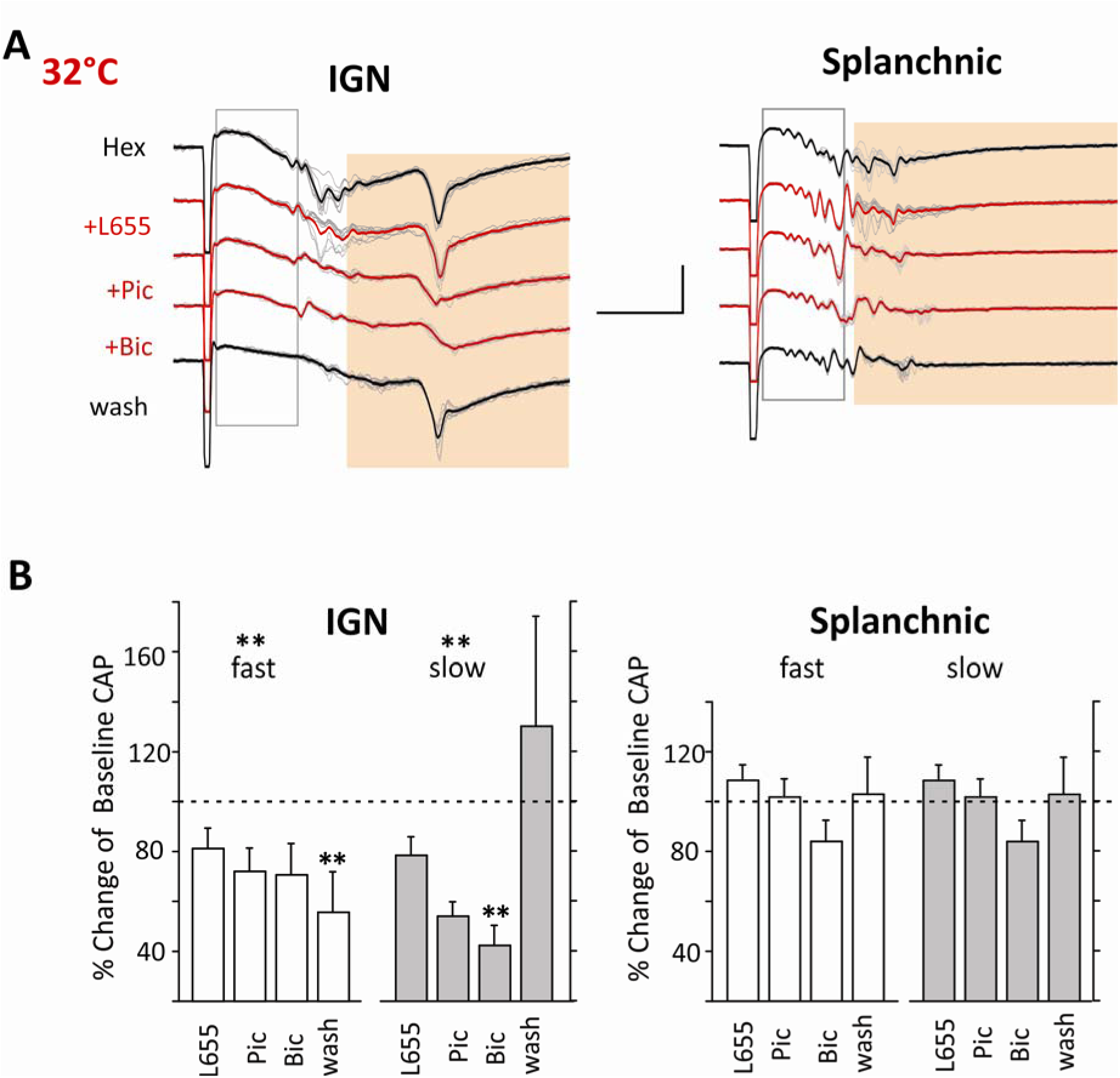
GABA_A_R antagonists increase conduction block in slow and branching SPN axons. [A]. Effect of GABA_A_R antagonists at 32°C. Shown are typical unfiltered raw traces of evoked responses, elicited through supramaximal stimulation using 200 µA/500 µs electrical impulses. Baseline response is in the presence of hexamethonium (Hex; 100 µM) to block synaptic transmission with subsequent application of L655,708 (L655; 50µM), picrotoxin (Pic; 50µM), and bicuculline (Bic; 10µM). The events have been high-pass filtered at 100 Hz. Displayed are 10 superimposed episodes for each condition, with their averages emphasized for clarity. **[B]** Significant block was seen only in slower axons in the IGN following application of Pic (p<0.05) and Bic (p<0.01). Results are displayed as mean % value of rectified integrals of fast and slow conducting responses relative to that obtained at baseline. Percentage value Statistical tests were undertaken using repeated measures ANOVA and Bonferroni t-test for all pairwise multiple comparisons (*, p<0.05; **, p<0.01; n=6). Scale bar is 20μV, 10ms.

The effects of the GABA_A_R positive allosteric modulator pregnanolone and of the GABA_A_R agonist muscimol were also tested (n=4). As with GABA_A_R antagonists there were no effects on splanchnic axon populations. In comparison, muscimol only acted on the fast IGN population with response reduction to 65.0±6.5% of control amplitude (p<0.01; n=4; not shown).

### Presynaptic acetylcholine (ACh) receptors are also a site for SPN axon conduction modulation

Following the blockade of ganglionic transmission with hexamethonium, we specifically assessed the cholinergic actions on presynaptic SPN axons by applying the cholinesterase inhibitor neostigmine and acetylcholine (ACh). Neostigmine (20µM) was used to prevent ACh degradation. Changes in evoked responses in the presence of neostigmine alone would be interpreted as due to increasing endogenously released ACh. Subsequently applied ACh (100µM) would be expected to have actions on receptors in relation to the dose applied. An example of evoked responses is shown in **Figure 3A**. Negative DC shifts were seen in 6/6 IGN and 5/6 splanchnic nerves following the application of neostigmine and ACh (**Fig. 3B_1_**). Mean voltage shifts were -348±63μV and -275±78μV, respectively. In all of these experiments (n=6/6), both IGN and splanchnic nerves exhibited an increase background spontaneous spiking activity after the application of neostigmine and ACh, consistent with the DC-shift being due to membrane depolarization (**Fig. 3B_2_**). ACh depressant actions were preferential to slow conducting axons with particularly strong block in IGN axons (to 6.5±2.1% of control, p<0.01) compared to splanchnic (to 44.6±15.6%;**, p<0.05; **Fig. 3C**). Note that spiking activity in the IGN could reflect increased activity in presynaptic SPN axons and/or postganglionic axons while that seen in the splanchnic nerve largely represents activity in preganglionic SPN axons.

**Figure 3.**
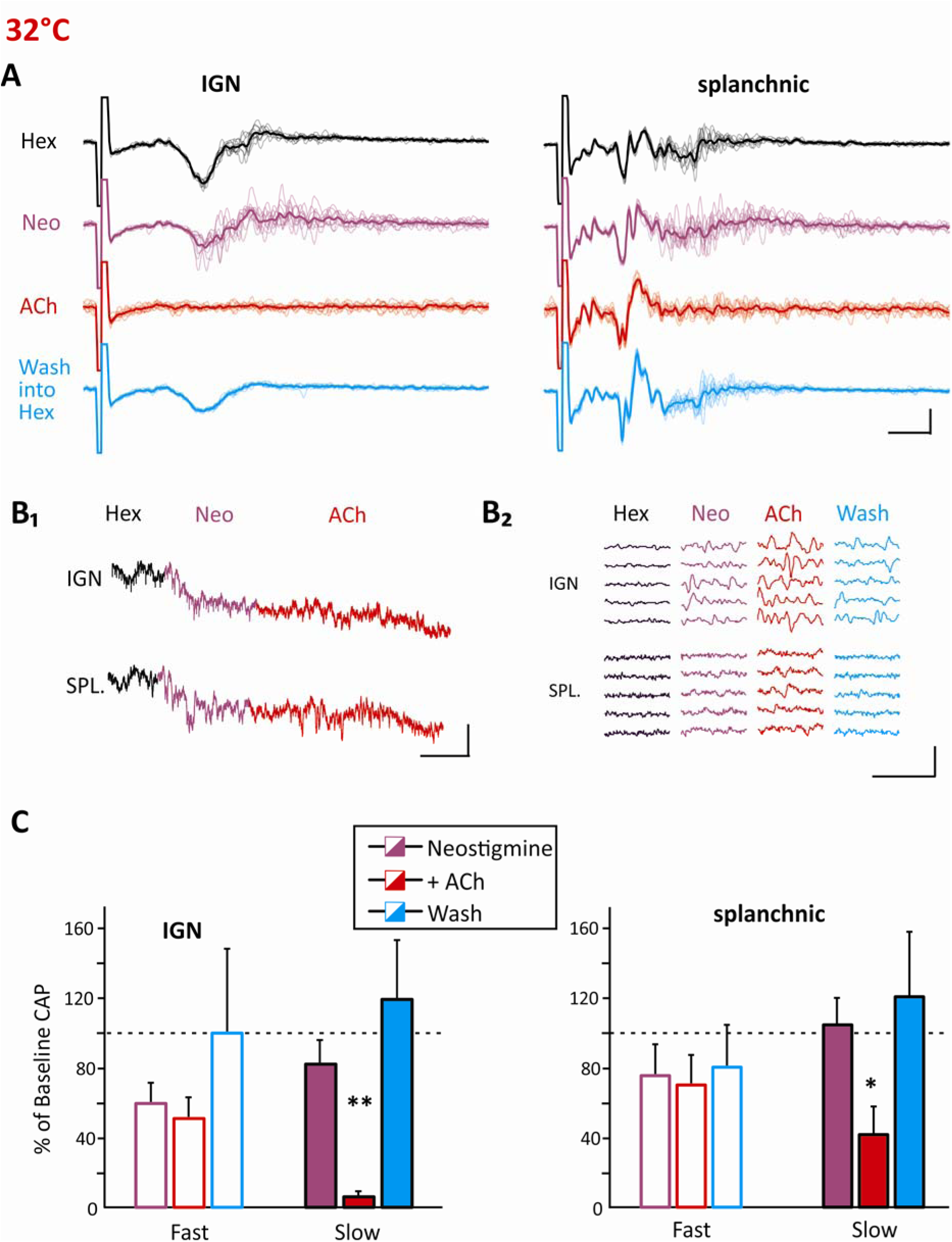
Evidence of presynaptic cholinergic modulation. [A]. Representative raw traces of electrically-evoked responses in IGN and splanchnic SPN axons, the effects of the cholinesterase inhibitor neostigmine (20µM) and ACh (100µM) application, and subsequent washout of applied drugs. Each trace is evoked by a 200 µA/500 µs electrical stimulus and has been high-pass filtered at 100 Hz for clarity. In all panels, drug applications are denoted by a change in the color of the trace. **[B_1_]** Example DC recordings showing clear increases in negativity following application of neostigmine and ACh. Clear negative DC shifts in IGN and splanchnic nerves were seen in 4 out of 6 experiments. Extracellular DC shifts after applied neostigmine and ACh reflect membrane depolarization. Recordings were undertaken coincident with stimulation of the ventral root every minute. **[B_2_]** Shown in an example of emergent spontaneous activity (particularly in IGN recordings) after applied neostigmine and ACh seen during DC shifts (demonstrated by a series of 5 stacked epochs taken 1 minute apart). Emergent spontaneous activity was seen in 6/6 preparations. **[C]** Applied ACh was associated with significant amplitude reductions in slow conducting axons: in the IGN (by 93.5±2.1%;**, p<0.01 post hoc Bonferroni t-test; RM ANOVA; ***, p<0.001)) and in Spl slow conducting axons (by 58.4±15.6%;**, p<0.05 post hoc Bonferroni t-test; RM ANOVA (***, p<0.01). N=6 for except for wash (where n=4). Scale bars are: A- 50μV, 5ms; B_1_- 100μV, 10min; B_2_- 10μV, 10ms.

The metabotropic ACh receptor antagonist atropine was tested on electrically-evoked responses at 32C (n=4) given known preganglionic M3 receptor expression (Alkaslasi et al., 2021). Atropine had selective depressant actions on the slower conducting splanchnic responses (70.6±11.0% of control; (p<0.05; Paired t-test).

### Pharmacological block of voltage-gated K^+^ channels

The shape and propagation of action potentials in neurons are significantly influenced by various post-spike voltage-gated K^+^ (K_V_) conductances. Among these, the fast-activated K_A_ channels accelerate spike repolarization to reduce duration. We tested actions of the K_A_ blocker 4-aminopyridine (4-AP). In compound action potential (**CAP**) population recordings of unmyelinated axons in the cerebellum and hippocampus, 4-AP leads to increases in CAP amplitude and duration (Palani et al., 2012), while reducing spike conduction failures (Pekala et al., 2016).

We first assessed dose-dependent changes in conduction by 4-AP at cumulative doses of 1, 5, 10, 50, and 100µM at 32°C (n=5). At this temperature the fastest-conducting myelinated axons in splanchnic could not be quantified due to its overlap with stimulus artifact. We used a four-parameter logistic model to obtain shared EC_50_ values across electrically- and optically-evoked responses in IGN and splanchnic SPNs separated into fast and slower conducting axons (**Fig. 4A**). Individual EC_50_ values (**Fig. 4A**) were generated in subsamples where quantifiable effects were seen at multiple doses (shown in **Fig. 4A**). Averaged over all groups with 4-AP-induced facilitation, the mean EC_50_ value was 41.0±3.7µM. As 50µM 4-AP is consistent with prior work using 4-AP for non-specific blockade of K_A_ channels, a dose of 50 or 100µM was used for subsequent studies on effects of spike conduction or recruitment.

**Figure 4.**
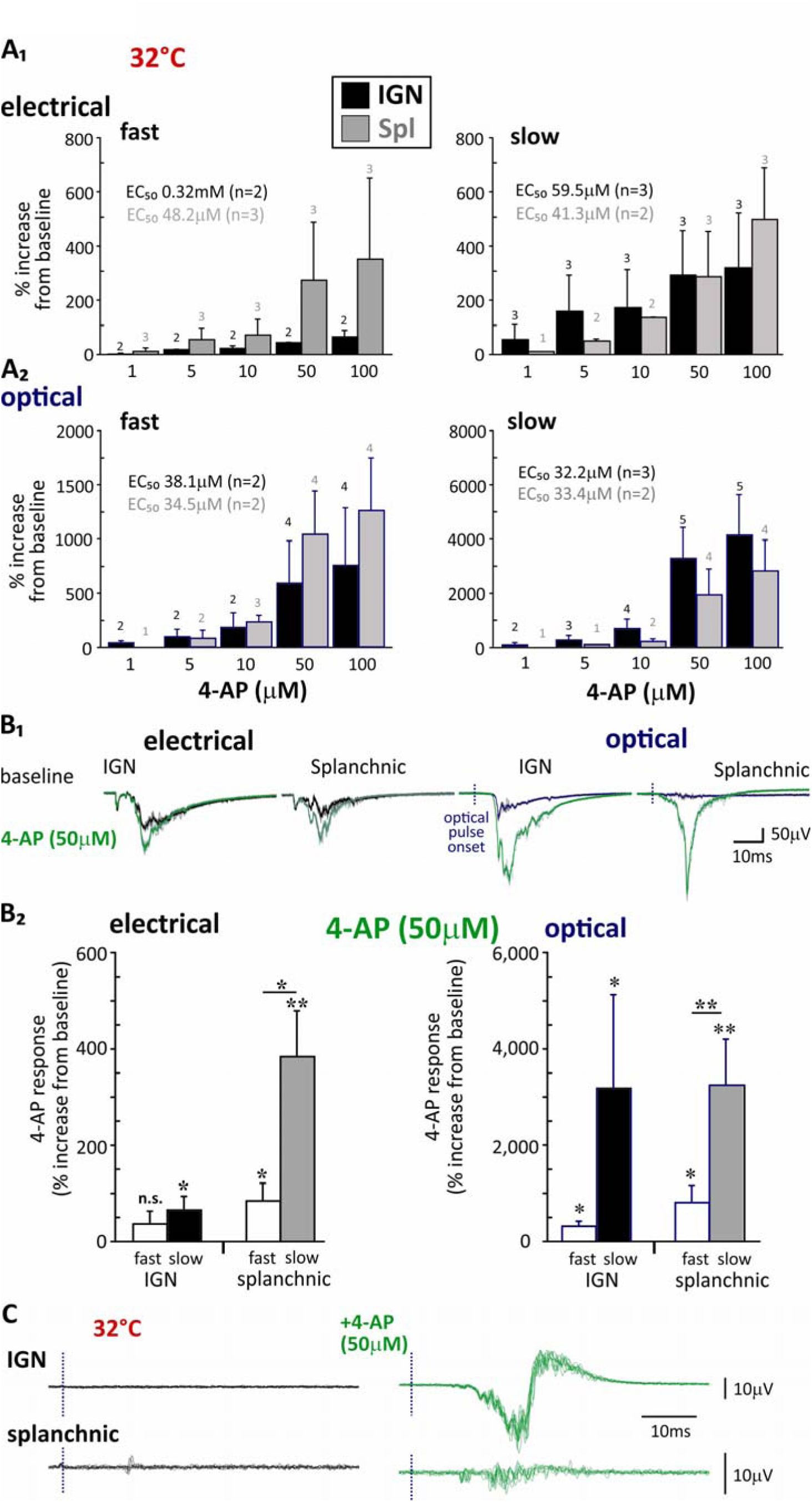
4-AP preferentially increases SPN conduction of slow-conducting SPNs. [A]. 4-AP dose-response effect on electrically and optically evoked responses in IGN and splanchnic nerves (n=5). Shown are dose-dependent increases in electrically (**A_1_**) and optically evoked responses (**A_2_**) separated into fast- and slower- conducting axons. Dose-response EC_50_ values (obtained from a subsample of mice) and observed response incidence at each dose (out of n=5) are shown in presented plots. Note that lower 4-AP doses were commonly without effect. **[B_1_]** Typical raw traces of evoked responses. The traces have been high-pass filtered at 25 Hz. Note the downward increase in amplitude following 4-AP. **[B_2_]** Quantification of the 4-AP response is plotted for fast- and slow-conducting SPNs. Similar response amplification is seen in IGN and splanchnic nerves. Note that greater actions are seen in slow-conducting SPNs, particularly those recruited optically. Paired t-tests were conducted to assess significance of amplitude of evoked responses between baseline and their percentage increase following application of 4-AP (n=6; *, p<0.05; **, p<0.01; error bars are SEM). Asterisks above bars represent significantly increased recruitment of slower over faster conducting axons. **[C]** Restoration of axonal conduction at 32°C in the presence of 4-AP. Example experiment without optically-recruited response at 32°C, application of 4-AP restored conduction in many SPN axons. The dotted line at beginning of raw recordings reflects timing of optical illumination. Scale bars are: B_1_- 50μV, 10ms; C- 10μV, 10ms.

Raw traces show the increased in evoked response following application of 50µM 4-AP (**Fig. 4B_1_**). 4-AP-induced increases in response amplitude were quantified following separation into fast and slow conducting components recruited by electrical or optical stimuli. Applied 4-AP led to significant increases in response magnitude for all populations except for fast-conducting responses within the IGN (n=6; **Fig. 4B_2_)**. Particularly impressive was the magnitude of response facilitation seen with optical stimuli; ∼5- to 10-fold greater than those seen electrically (p<0.05 for all groups except fast splanchnic which is p=0.08; **Table 4**). The much greater 4-AP induced increase in evoked responses optically is consistent with their preferential temperature dependent conduction failures shown (**Fig. 1C)** and likely accounts for the dramatic potentiation seen relative to electrical after 4-AP. For example, application of 4-AP successfully restored conduction in cases where optically-recruited responses were either minimally observable (n=5/6) or completely absent (n=1/6) in the IGN (**Fig. 4C**).

**Table 4.** 4-AP induced increase in response magnitude comparison between fast and slow in electrically and optically evoked responses. Comparing the amplitude of electrical, optical, versus co-stimulation evoked responses from the same ventral root. Values represent % increase relative to baseline response magnitude in IGN and splanchnic nerve recordings (n=6). Paired t-tests assess significance. (*, p<0.05; **, p<0.01)

| % increase | Electrical | Optical | E vs. O |
| --- | --- | --- | --- |
| IGN - fast | 38±27 (p=0.2) | 326±105* | * |
| IGN - slow | 67±28* | 3,195±1,948** | * |
| Spl - fast | 86±36* | 820±357* | (p=0.08) |
| Spl - slow | 385±96* | 3,261±953** | * |

In concert with our observation that overall electrical stimulation recruited population conduction is reduced by ∼50% when temperature is elevated from 22°-36°C(Halder and Hochman, 2026), the present data suggest that activation of activation of K_A_ K^+^ channels plays a role in conduction failures and that suppression of their activity with 4-AP is capable of dramatic response potentiation. This finding underscores the significant role of post-spike voltage-gated K^+^ channels in modulating axonal conduction.

### Tetraethylammonium (TEA) has unexpected inhibitory actions and acts independent of 4-AP

Like 4-AP, the nonspecific voltage-gated K^+^ channel blocker TEA (Kirchhoff et al., 1992) has been shown to reduce conduction failures in unmyelinated cerebellar granule axons, likely by an increased spike duration (Pekala et al., 2016). We compared the effects of TEA (1 mM) to that of 4-AP on evoked responses as shown in example recordings (**Fig. 5A)**. In contrast to expectations, TEA had no effect on response amplitude of faster-conducting IGN axons or on faster- or slower-conducting splanchnic axons. Furthermore, it selectively depressed rather than facilitated the amplitude of slower-conducting axons of the IGN by 81.1% (p<0.01; **Fig. 5B**). In comparison, subsequent application of 4-AP invariably led to large response increases in all populations as was observed in the absence of TEA (**Fig. 5A, B**). These results demonstrate that TEA and 4-AP do not act on common sites.

**Figure 5.**
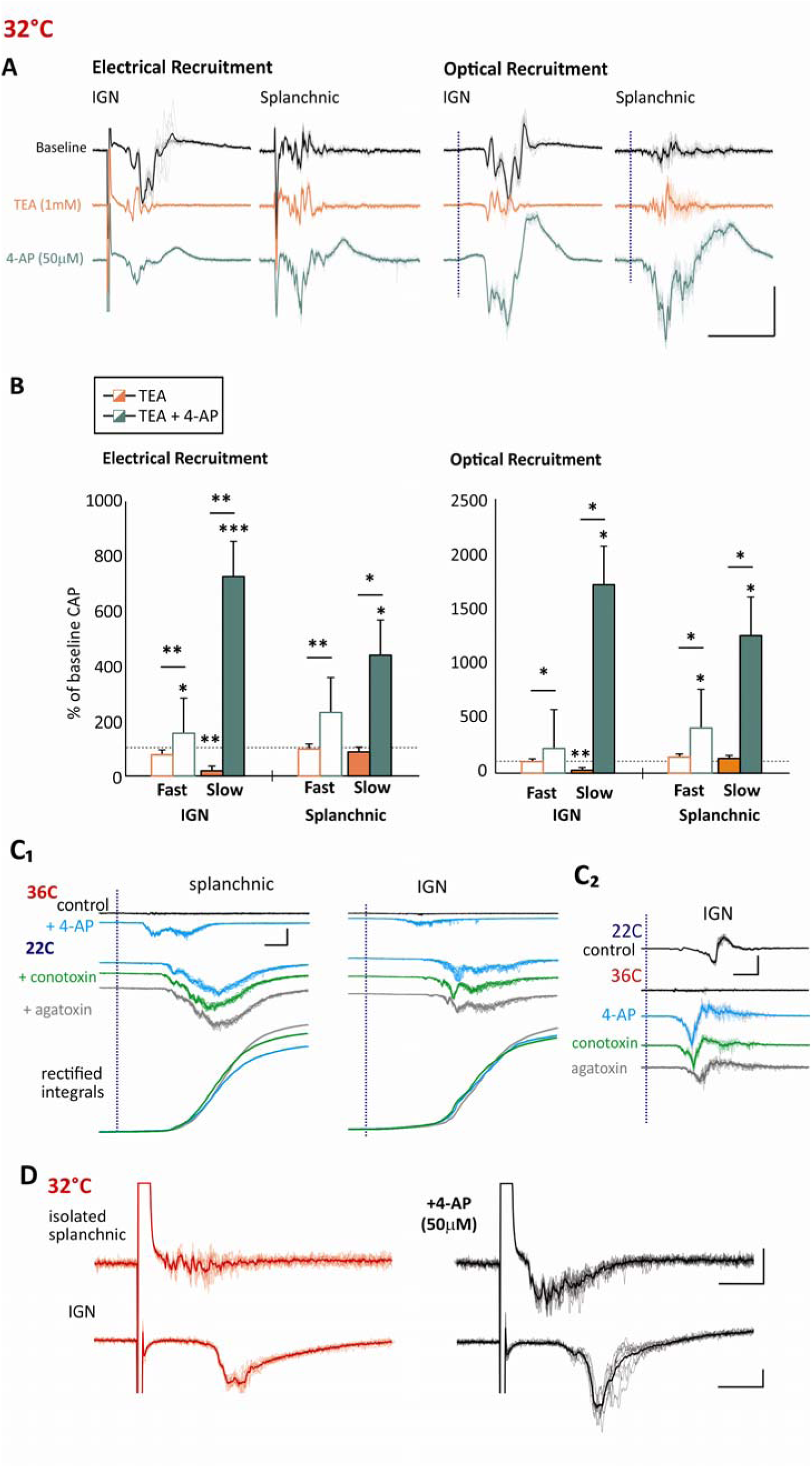
Differential actions of K^+^ channel blockers TEA and 4-AP. [A]. Shown are typical raw traces of evoked responses, elicited through supramaximal stimulation using 200 µA/500 µs electrical impulses and 6.6mW/mm^2^ 20ms optical impulses. The traces have been high-pass filtered at 100 Hz. **[B]** Quantification of the evoked responses for each drug treatment is plotted for fast and slow conducting SPNs. 4-AP response was significant for all fiber types compared to hexamethonium baselines except for fast-conducting splanchnic axons. TEA exhibited a significant decrease from baseline in electrically and optically recruited slow-conducting IGN axons (p<0.01) but was otherwise no different from baseline. In all cases, the 4-AP response was significantly greater than the TEA response. Experiments were conducted at 32° C. Repeated measures ANOVA were employed to determine the significance of drug application effects, followed by a post hoc T-test for a comparative analysis between TEA and 4-AP (n=4; *, p<0.05; **, p<0.01; ***, p<0.001; error bars are SEM). **[C]** Ca_V_ channels have limited effect on 4-AP-induced facilitation. Shown are two panels that assess the effects of Ca_V_ antagonists on optically evoked responses in the presence of 4-AP at 36 (left; 100 Hz high-pass filtered) and 22°C (right; unfiltered). Panel at right also includes rectified-integrated averages of responses to more clearly demonstrate lack of drug actions. Drug doses are: 4-AP (100μM), ω-conotoxin GVIA (50nM), and ω-agatoxin IVA (50nM). **[D]** 4-AP induced potentiated response is seen in the isolated splanchnic nerve and is comparable to that seen in the T12 IGN. Shown are electrically-evoked responses at 32°C. Scale bars are: A- 50μV, 10ms; C- 400μV, 10ms; D-50μV, 10ms.

As TEA blocks ganglionic transmission (Shand, 1965), TEA-generated depression of the slower arriving IGN volley could be explained by block of synaptic postganglionic spiking responses. This assumes the preincubated ganglionic blocker hexamethonium (100 μM) was insufficient to block postganglionic spiking. To test whether 4-AP facilitatory actions on slower conducting responses in the IGN included recruitment of postganglionic spiking by surmounting hexamethonium block of nicotinic ACh receptors, we examined the effects of 4-AP after block of voltage-gated Ca^2+^ (Ca_V_) channels (**Fig. 5C)**.

Preganglionic synaptic transmission is thought to be primarily via N but also P-type Ca_V_ channels (Ireland et al., 1999). 4-AP facilitatory may include the selective actions on the Cacnb3 (β3 unit) of the N-type Ca_V_2.2 Ca^2+^ channel (Wu et al., 2009), whose expression is limited to 2 of 15 thoracic SPN subpopulations (Alkaslasi et al., 2021). The N-type blocker ω-conotoxin GVIA (50nM) partially depressed 4-AP facilitation in 1/3 mice. The P/Q-type blocker ω-agatoxin IVA (50nM) led to a rightward shift consistent with reduction in conduction velocity but had little effect overall (n=2 mice). Thus, the potentiation produced by 4-AP is not due to the emergence of postsynaptic postganglionic responses in CAP recordings.

We next assessed the effects of 4-AP on electrically evoked responses in the completely isolated splanchnic nerve (n=3). In all occasions 4-AP led to a clear response facilitation, and more importantly, the actions had comparable negative polarity appearance to that seen with intact IGN and splanchnic nerves, further ruling out synaptically-mediated effects **(Fig. 5D).**

### Role of K_2_P leak channels in the temperature-dependent in block of spike conduction

DC recordings were undertaken in IGN and splanchnic nerves to examine the shifts in membrane polarization associated with temperature changes between 22°C and 32°C (**Fig. 6A_1_**). Extracellular negative or positive shifts in polarity should reflect depolarizing or hyperpolarizing shifts in membrane potential, respectively. Elevating temperature consistently led to positive DC shift in polarity in both splanchnic and IGN nerves indicative of membrane hyperpolarization (2.8±0.3 and 2.9±0.3mV, respectively; n=5/5). Conversely, lowering temperature from 32°C to 22°C always produced a corresponding negative DC shift reflective of membrane depolarization (−3.1±0.8 and -2.7±0.3mV, respectively; n=3/3). As the splanchnic nerve likely contains few postganglionic axons (Baron et al., 1985), we interpret the comparable DC polarizing actions in the IGN, which also contains postganglionic axons, to be driven by SPN axons.

**Figure 6.**
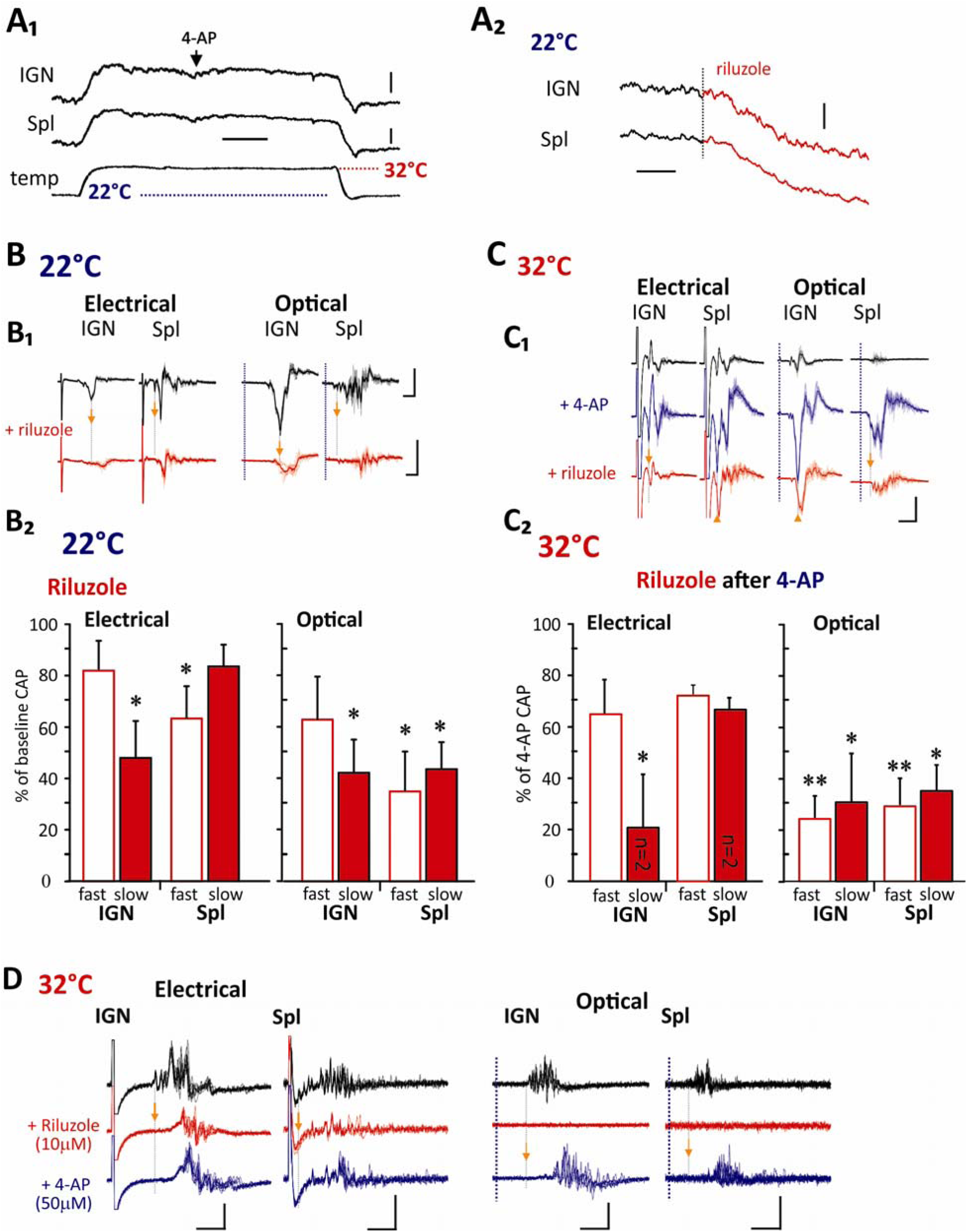
Effect of riluzole at 22°C and at 32°C subsequent to K^+^ channel block with 4-AP. [A]. DC shifts. **[A_1_]** Bath temperature increases from 22° to 32° lead to a reversible depolarizing DC shift. **[A_2_]** Riluzole also consistently leads to hyperpolarizing DC shifts. **[B]** Effects of riluzole on evoked responses. **[B_1_]** Shown are typical evoked responses in SPNs to riluzole (50µM; 100 Hz high-pass filtered). **[B_2_]** The graph shows the depressant effects of riluzole (50µM) on the rectified integral of the maximal CAP (n=4). **[C]** Riluzole depressant actions also occur in the presence of 4-AP. **[C_1_]** Shown are typical evoked responses to 4-AP (50µM) and riluzole (10µM). **[C_2_]** The graph shows the influence of riluzole (concentration range 0.5-50µM) on the CAP induced by 4-AP (n=3 except where stated). **[D]** Riluzole was administered before the application of 4-AP which reduced electrically evoked CAP while eliminating the optically evoked CAP. However, the subsequent introduction of 4-AP resulted in an increase in electrically evoked SPNs and successfully restored the optically evoked SPNs (n=4). The orange arrows with grey dotted lines highlight the delay in CAP conduction after applied riluzole. All statistics were undertaken with paired t-tests (*, p<0.05; **, p<0.01). Responses were 100 Hz high-pass filtered. Scale bars are; A_1_- 2mV, 10min; A_2_- 400μV, 5min; B_1_- 250μV, 10ms; C_1_-10μV, 10ms; D- 25μV, 10ms (IGN) and 2.5μV, 10ms (Spl).

### Potassium leak channel activators riluzole and arachidonic acid block axon conduction

Temperature increase driven changes in membrane potential are consistent with activation of temperature-sensitive two-pore K^+^ leak (**K_2_P**) channels with expected increase in membrane leak conductance. Consequently, their activation may play a significant role in observed conduction failures at increased temperature(Halder and Hochman, 2026). The polyunsaturated fatty acid arachidonic acid (**AA**) and riluzole can activate the TREK-1 K_2_P channel (Maingret et al., 2000; Cadaveira-Mosquera et al., 2011). In comparison, traditional voltage gated K^+^ channel blockers TEA and 4-AP are without effect on K2P channels (Enyedi and Czirják, 2010),

We first assessed the effect of riluzole at 22°C where the temperature sensitive TREK-1 channel should be minimally activated (**Fig. 6A_2_, B**; n=4). Riluzole consistently generated a small DC shift in both GN and splanchnic nerves (−668±238 and -681±243μV, respectively). Regarding electrically-evoked responses, riluzole preferentially decreased activity in slower-conducting IGN axons (to 48.1±14.1%; p<0.05) and faster-conducting SPN axons (to 63.2±12.5% of baseline; p<0.05). Regarding optically evoked responses, decreases were seen in only slower-conducting IGN axons (to 42.3±12.7%; p<0.05) while depression was seen in both faster- and slower-conducting splanchnic axonal populations (to 35.1±15.4%; p<0.05 and 43.7±10.4% of baseline; p<0.05, respectively).

As riluzole also activates K_A_ currents, to better focus riluzole actions on K_2_P leak channels we blocked K_A_ with 4-AP before applying varying concentrations of riluzole (0.5-50µM) (Duprat et al., 2000) (**Fig. 6C**; n=3). These studies were undertaken at 32°C to further activate the temperature sensitive TREK-1 channels. Regarding electrically-evoked responses, riluzole only decreased activity in slower-conducting IGN axons (to 21.2±21.2% of baseline; p<0.05). Regarding optically evoked responses, decreases were seen in both faster- and slower-conducting axons in both nerves. For IGN fast and slow populations, reductions were to 23.8±11.7% (p<0.01) and 30.1±19.0% (p<0.05) of baseline, respectively. For splanchnic fast and slow populations reductions were to 28.3±13.1% (p<0.01) and 35.4±10.8% (p<0.05) of baseline, respectively. Note that riluzole actions also included a reduction in conduction velocity (**Fig. 6B_1_ & C_1_** – shaded bars), consistent with expected actions on K_2_P leak channels in decreasing membrane resistivity.

We previously showed that increased temperature can lead to complete conduction block of optically-evoked responses, presumed to be in part via activation of K_2_P leak channels. At 32°C, we applied riluzole to see if there was additional depression of responses (n=4). In 2/4 experiment riluzole (50µM) completely abolished optically recruited responses (**Fig. 6D**). Subsequent administration of 4-AP (50µM) potentiated electrical (4/4) and optical responses (2/4 cases), but only restored 1/2 responses abolished by riluzole. Overall findings indicate a preferential depression of optically evoked responses to riluzole treatment, particularly at 32°C. That block of optically evoked responses was associated with conduction delay in electrically-evoked responses, support loss of conduction due to increased membrane leak.

The effects of arachidonic acid was assessed at 1, 5 and 10 µM (n=5). Compared to riluzole, arachidonic acid (10µM) actions were more selective with slight DC shifts in 4/5 mice (means of 81±31 and 89±32 μV in IGN and splanchnic recordings, respectively; **Fig. 7A)**. At 22°C, arachidonic acid preferentially led to a reduction in the amplitude of slow conducting IGN electrical and optical responses by 46.9±6.9% (p<0.001) and 42.9±8.2% (p<0.01), respectively; **Fig. 7B,C**). Significant effects were also seen in slow conducting responses in the IGN at 5µM (by 27.9±9.4% [p<0.05] and 28.5±7.1% [p<0.01], respectively) and 1 µM in IGN optically evoked responses (by 19.8±7.4%; p<0.05).

**Figure 7.**
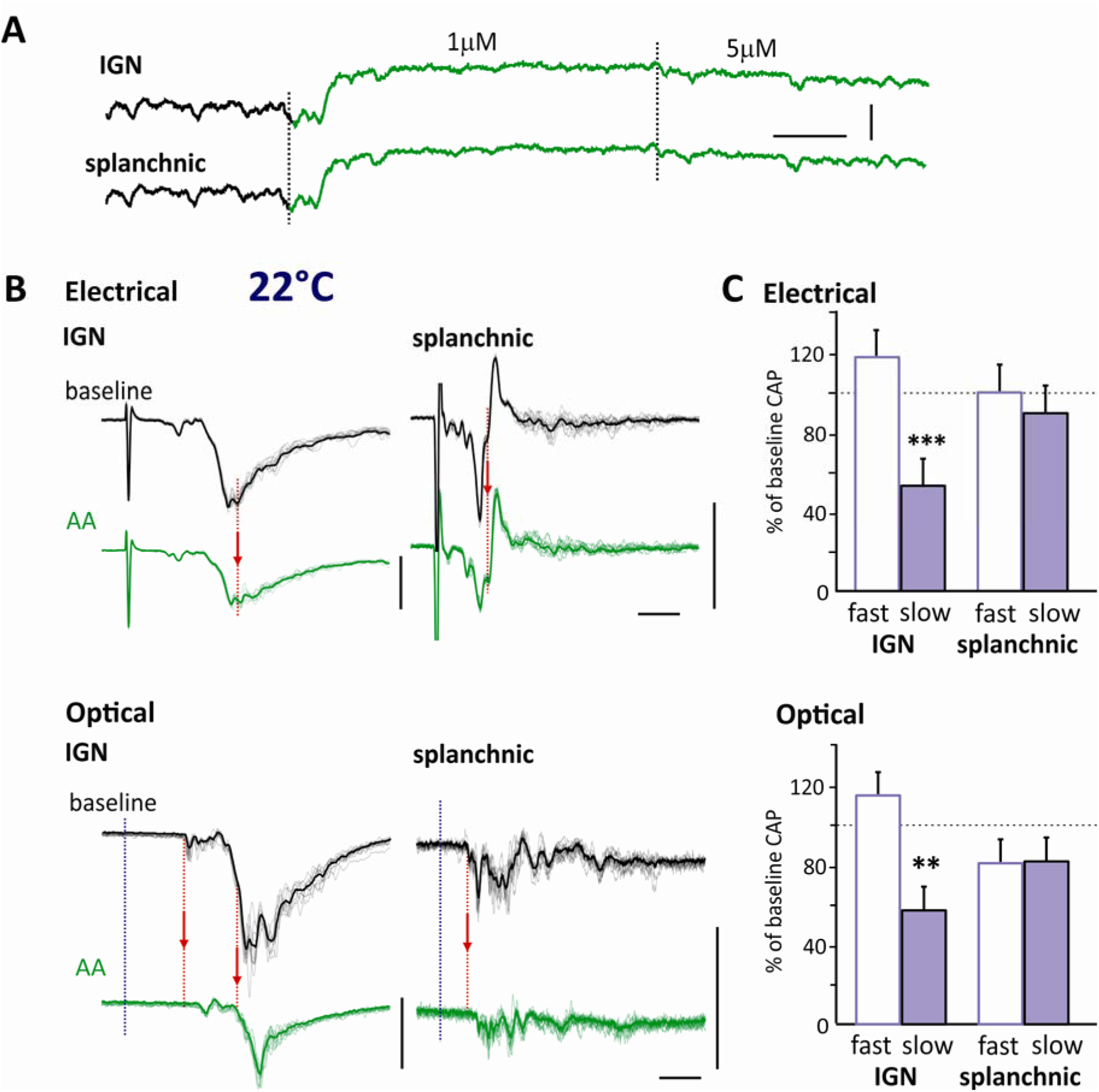
Arachidonic acid contributes to blocking slow-conducting SPN fibers recruited optically and electrically. Experiments were performed at 22°C. **[A]** Example DC shift after bath applied arachidonic acid at doses shown. **[B]** Example evoked responses in SPNs, triggered by supramaximal electrical or optical impulses before and after application of arachidonic acid (AA) at 10 µM. Traces are presented as 10 superimposed unfiltered raw traces with overlaid average. Note that AA led to a conduction velocity slowing in some identifiable components of the CAP volley (red arrows). Traces are presented as 10 superimposed unfiltered raw traces with overlaid average. **[C]** AA (10µM) preferentially reduced CAP responses in slow-conducting IGN axons evoked by electrical and optical stimulation. Statistical analysis was conducted using paired t-tests to determine the significance (n=5; **, p<0.01; ***, p<0.001). Scale bars are [A] 100μV, 5 min; [B] Electrical- 50μV, 10ms; Optical- 25μV, 10ms.

## DISCUSSION

The present study demonstrates that conduction along thoracic sympathetic preganglionic neuron (SPN) axons is highly dynamic and strongly regulated by ion channel activity, neuromodulatory receptor activation, and temperature-dependent changes in membrane conductance. Rather than functioning as passive transmission cables, SPN axons are prone to conduction failure. In particular, the slow-conducting unmyelinated fibers traversing the interganglionic nerve (IGN), appear to operate near the threshold for reliable propagation and can undergo profound facilitation or failure depending on physiological state. Together with our companion studies (Halder and Hochman, 2026; Halder et al., 2026), these findings establish that spike propagation in thoracic SPN axons is probabilistic rather than invariant and that conduction reliability can be dynamically regulated by physiological and pharmacological mechanisms. These findings further raise the possibility that variable conduction reliability contributes to previous observations that a substantial fraction of anatomically identified SPNs exhibit little ongoing or reflex-related activity (Jänig, 2022). Slow branching axon (IGN) responses were consistently more vulnerable than unbranching (splanchnic nerve), supporting the idea that branch-point geometry is a major determinant of propagation safety factor. Similar branch-point failures have been described in sensory systems where propagation reliability can be regulated dynamically by membrane polarization and GABA_A_ receptor activity (Wall, 1994; Wall and McMahon, 1994; Wall, 1995; Lucas-Osma et al., 2018; Hari et al., 2022). The present findings extend this concept to sympathetic pathways and suggest that regulation of axonal conduction may serve as a distributed mechanism for controlling autonomic gain. A summary of these findings is provided in **Figure 8**.

**Figure 8.**
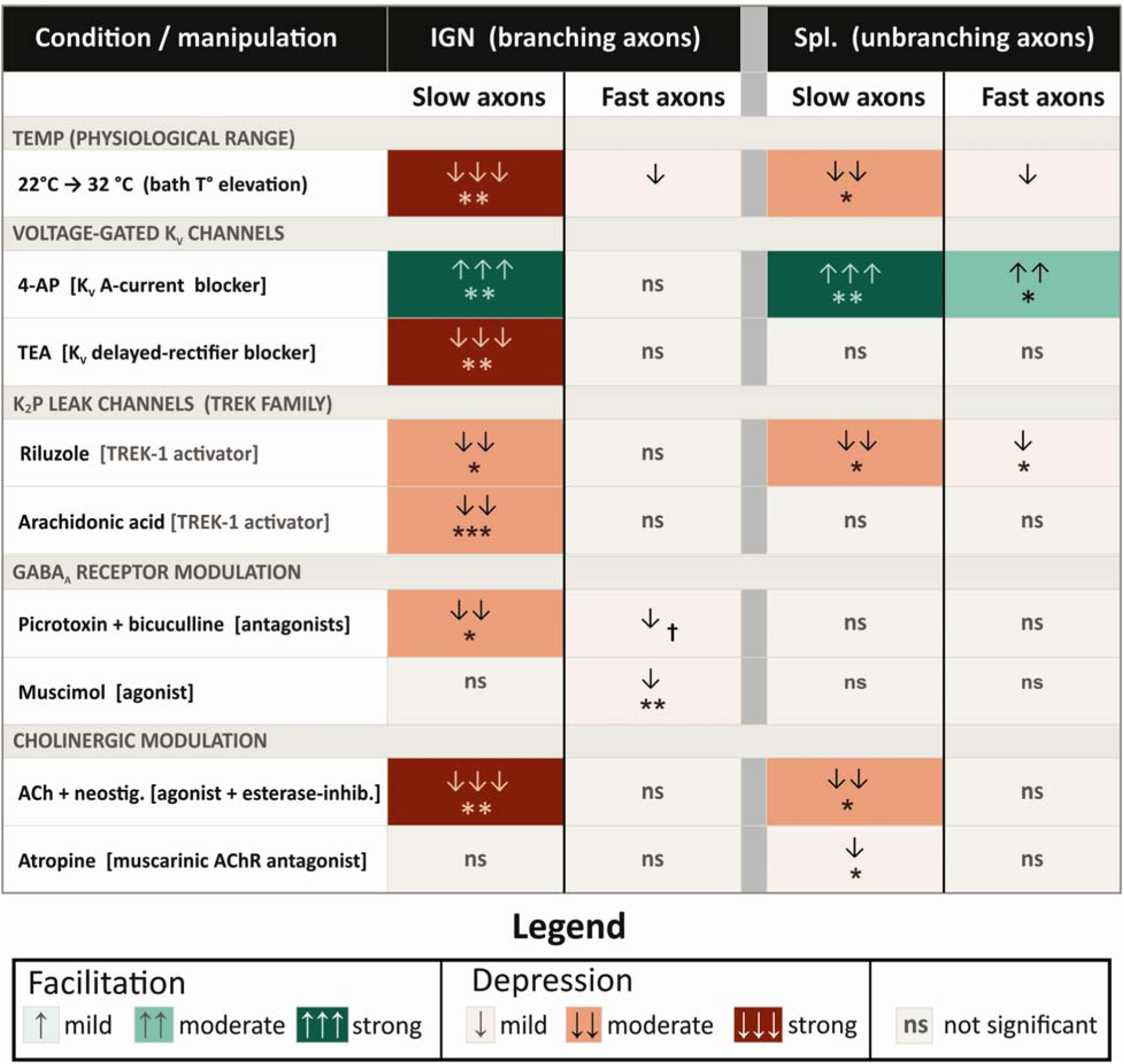
Pharmacological fingerprint of SPN axon subpopulations. Direction and magnitude of conduction modulation across branching (IGN) and unbranching (splanchnic) SPN axons by each pharmacological condition. Cell color = direction and magnitude of effect on CAP area (rectified integral). Magnitude tiers: ↓ = 20–40% reduction, ↓↓ = 40–60%, ↓↓↓ >60%; ↑↑ = 2–5× increase, ↑↑↑ >5×. Significance: *, p<0.05; **, p<0.01; ***, p<0.001; † = effect at washout only. All experiments at 32°C except riluzole and arachidonic acid (22°C). Electrical and optical responses pooled were concordant.

### Ion channel contributions

(a) <u>K2P leak channels.</u> Propagation reliability was highly temperature-sensitive, particularly within optically recruited and slow-conducting axonal populations. Elevating bath temperature from 22°C to 32°C consistently produced extracellular DC shifts indicative of membrane hyperpolarization and strongly reduced conduction, in some cases abolishing optically recruited responses entirely. These findings strongly implicate activation of potassium leak conductances. TREK-family K2P channels are attractive candidates because they are temperature sensitive, expressed in subsets of SPNs, and would be expected to reduce membrane resistance and impair current transfer at branch points (Maingret et al., 1999; Maingret et al., 2000; Alkaslasi et al., 2021; Blum et al., 2021).

Consistent with this interpretation, the TREK activators riluzole and arachidonic acid preferentially depressed slow-conducting IGN responses, and slowed conduction velocity. Although the pharmacological profile is consistent with activation of temperature-sensitive K2P channels, particularly TREK-family K_2_P channels, these findings should be interpreted cautiously because neither riluzole (Debono et al., 1993; He et al., 2002; Bellingham, 2013) nor arachidonic acid (actions on multiple transduction cascades) is selective for TREK-family channels. Thus, the present data support K_2_P channel involvement but do not definitively identify the molecular species responsible for the observed changes in propagation reliability.

(b) Voltage-gated K^+^ channels. 4-AP robustly facilitated propagation, recruited previously silent axons, and restored temperature-sensitive conduction failures, in some cases re-establishing responses that were nearly absent at 32°C. Together, these findings indicate that many SPN axons normally operate close to propagation threshold and that modest changes in K^+^ conductance can profoundly alter conduction reliability.

The facilitatory actions of 4-AP are most consistent with blockade of A-type K^+^ conductances. Kv4-family channels are strongly expressed in SPNs (Alkaslasi et al., 2021), relatively insensitive to TEA, and known to accelerate spike repolarization and shorten action potential duration (Jerng et al., 2004). Previous studies demonstrated that I_A_-like axonal conductances can regulate propagation directly (Debanne et al., 1997), while 4-AP restores conduction in demyelinated and temperature-sensitive mammalian axons (Sherratt et al., 1980; Bostock et al., 1981). Spike broadening caused by suppression of I_A_ conductances would increase axial current transfer to downstream branch segments and thereby reduce branch-point failure.

The contrasting actions of 4-AP and TEA are particularly informative. Although both drugs are commonly classified as K^+^ channel blockers, they produced strikingly different effects on SPN conduction reliability. Whereas 4-AP facilitated propagation, TEA selectively depressed slow-conducting IGN responses. These observations indicate that the conductances governing propagation security in SPN axons are unlikely to be dominated by classical delayed rectifier currents preferentially blocked by TEA. Instead, the data supports a prominent role for 4-AP-sensitive conductances, presumably A-type channels, in determining propagation safety factor. Such effects are consistent with improved branch-point propagation.

TEA has previously been reported as a blocker of ganglionic transmission (Shand, 1965) and presumed to be via <u>postsynaptic</u> depolarization block of spiking (Paton and Perry, 1953) where block of multiple voltage-gated K^+^ channels leads to a progressive depolarization with subsequent maintained inactivation of voltage-gated Na^+^ channels. It is therefore possible that depressant actions of TEA on slow IGN axons may be via inadequate block of synaptic transmission with hexamethonium (100µM) in this subpopulation of axons. However, the persistence of 4-AP facilitation following blockade of N-type and P/Q-type Ca^2+^ channels, together with similar effects observed in isolated splanchnic nerves lacking postganglionic neurons, argues against a major contribution from ganglionic transmission. A more parsimonious interpretation is that TEA directly reduces propagation reliability in vulnerable SPN axons, potentially through depolarization-dependent sodium channel inactivation resulting from block of resting potassium conductances.

(c) Voltage-gated Ca^2+^ channels. The Ca^2+^ channel antagonist experiments further support modulatory actions via directs actions on SPN axons. Persistence of 4-AP facilitation in isolated splanchnic nerves and after blockade of N-type and P/Q-type Ca^2+^ channels argues against recruitment of postganglionic activity as the primary explanation for 4-AP-induced potentiation.

Although classically associated with neurotransmitter release, voltage-gated Ca² channels localized to axons can directly influence spiking independently of synaptic transmission (Bender and Trussell, 2009). N-type channels contribute to action-potential-associated Ca^2+^ influx regulating Ca^2+^-activated K^+^ conductances and afterhyperpolarization (Sah and McLachlan, 1995; Wikström and Manira, 1998; Ireland et al., 1999), while P/Q-type channels are tightly coupled to Ca^2+^-activated K^+^ currents shaping repolarization dynamics (Womack et al., 2004).

### Receptor mediated modulation of conduction

The present findings also identify neuromodulatory receptor systems as important regulators of propagation reliability. Antagonism of GABA_A_ receptors selectively depressed slow-conducting IGN axons, supporting a role for tonic GABA_A_ receptor activity in facilitating propagation. These findings parallel recent demonstrations that extrasynaptic α5-containing GABA_A_ receptors near branch points of sensory afferents produce tonic depolarization facilitating spike transmission (Lucas-Osma et al., 2018; Hari et al., 2022). Transcriptomic studies indicate that SPNs express α5-containing GABA_A_ receptor subunits (Wang et al., 2008; Alkaslasi et al., 2021), raising the possibility that similar branch-point mechanisms operate in sympathetic pathways. These findings extend axonal GABA_A_-mediated regulation of spike propagation from sensory afferents to autonomic pathways and are consistent with modulation occurring at sites of low propagation safety factor.

The directionality of GABA_A_ receptor actions depends critically on chloride equilibrium potential. Limited evidence for KCC2 expression in axons (Williams et al., 1999) suggests chloride equilibrium may remain relatively depolarized in these compartments, permitting GABA_A_ receptor activation to enhance rather than suppress excitability. Interestingly, muscimol preferentially depressed faster IGN axons rather than slow-conducting populations. This finding was unexpected and suggests that chloride homeostasis, receptor density, or receptor subtype composition may differ substantially across SPN axonal populations. Alternatively, exogenous receptor activation may engage mechanisms distinct from those associated with tonic endogenous GABA_A_ receptor activity.

Cholinergic modulation also strongly influenced propagation reliability. Following blockade of ganglionic transmission with hexamethonium, acetylcholine i(n the presence of the cholinesterase inhibitor neostigmine) generated DC shifts and spontaneous activity consistent with membrane depolarization while selectively suppressing slow responses – with near-complete conduction block in branching IGN axons. The combination of membrane depolarization, increased spontaneous activity, and reduced evoked propagation is consistent with depolarization-dependent Na^+^ channel inactivation. Under this interpretation, modest depolarization initially increases excitability whereas sustained depolarization reduces the availability of voltage-gated Na^+^ channels required for reliable spike propagation. Such a mechanism would be expected to disproportionately affect slow-conducting branching axons already operating near the limits of conduction safety. Both nicotinic and muscarinic receptors are expressed in SPNs (Alkaslasi et al., 2021; Blum et al., 2021), although the receptor populations responsible for these effects remain uncertain.

### Differences between electrical and optical recruitment

The present data also reveals important differences between electrical and optical recruitment. Simultaneous electrical and optical stimulation produced responses larger than either stimulus alone, indicating only partial overlap in recruited populations. Optical stimulation additionally exhibited much greater sensitivity to temperature-dependent failure and dramatically larger facilitation by 4-AP. These findings suggest that different SPN subpopulations possess substantially different propagation safety factors and pharmacological sensitivities.

### Limitations

Several limitations should be considered. First, the use of population compound action potential recordings does not permit direct identification of individual axons or precise localization of conduction failures to specific branch points. Second, the pharmacological agents employed are not fully selective for individual channel subtypes, limiting mechanistic attribution at the molecular level. Third, although ventral root afferents have been anatomically described (Coggeshall et al., 1977; Coggeshall, 1979), the contribution of their recruitment via electrical stimulation is likely minimal given the strong concordance between electrical and optical responses as optogenetic stimulation via ChAT-Cre/ChR2 recruit cholinergic axons. Finally, these experiments were performed under controlled conditions *ex vivo* and are unlikely to fully reflect ongoing neuromodulatory influences present *in vivo*.

### Perspective and impact

The present findings establish axonal conduction reliability as a major site of sympathetic gain control. Branching SPN axons represent a critical bottleneck where relatively small changes in ion channel activity, temperature, or neuromodulatory tone can produce large changes in sympathetic output. This framework substantially expands the traditional view that sympathetic regulation is governed primarily by synaptic integration within the spinal cord and ganglia. Instead, the axon itself emerges as an active computational element regulating amplification and distribution of sympathetic drive. Because conduction reliability can be modified pharmacologically, these mechanisms may represent clinically relevant targets in disorders characterized by altered autonomic output.

## Acknowledgements

This research was funded by NIH NS121850, NS102871

